# Patient DNA cross-reactivity of the CDC SARS-CoV-2 extraction control leads to an inherent potential for false negative results

**DOI:** 10.1101/2020.05.13.094839

**Authors:** Adam P. Rosebrock

## Abstract

Testing for RNA viruses such as SARS-CoV-2 requires careful handling of inherently labile RNA during sample collection, clinical processing, and molecular analysis. Tests must include fail-safe controls that affirmatively report the presence of intact RNA and demonstrate success of all steps of the assay. A result of “no virus signal” is insufficient for clinical interpretation: controls must also say “The reaction worked as intended and would have found virus if present.” Unfortunately, a widely used test specified by the US Centers for Disease Control and Prevention (CDC) incorporates a control that does not perform as intended and claimed. Detecting SARS-CoV-2 with this assay requires both intact RNA and successful reverse transcription. The CDC-specified control does not require either of these, due to its inability to differentiate human genomic DNA from reverse-transcribed RNA. Patient DNA is copurified from nasopharyngeal swabs during clinically-approved RNA extraction and is sufficient to return an “extraction control success” signal using the CDC design. As such, this assay fails-unsafe: truly positive patient samples return a false-negative result of “no virus detected, control succeeded” following any of several readily-encountered mishaps. This problem affects tens-of-millions of patients worth of shipped assays, but many of these flawed reagents have not yet been used. There is an opportunity to improve this important diagnostic tool. As demonstrated here, a re-designed transcript-specific control correctly monitors sample collection, extraction, reverse transcription, and qPCR detection. This approach can be rapidly implemented and will help reduce truly positive patients from being incorrectly given the all-clear.

**One Sentence Summary:** A widely-used COVID-19 diagnostic is mis-designed and generates false-negative results, dangerously confusing “No” with “Don’t know” – but it’s fixable

## Introduction

Molecular diagnostic testing has become (socially-distanced) dinner table conversation during the COVID-19 pandemic. Reverse-transcription coupled quantitative polymerase chain reaction (RT-qPCR) has been widely deployed to measure viral RNA genomes present in patient samples. SARS-CoV-2 RT-qPCR assays can detect virus from early stages of infection *(1)*. Measuring viral RNA informs as to presence of virus at the time of sampling and can be used as a measure of potential infectivity, complementing serological testing that can only indicate past infection. During the first months of the COVID-19 pandemic, testing has focused on enumerating positive patients in the context of triage and epidemiological monitoring. As the world begins to reopen, the focus of molecular testing will shift to *clearing* asymptomatic individuals based on absence of detectable virus.

The US Centers for Disease Control and Prevention (CDC) have specified and given emergency use authorization (EUA) for a SARS-CoV-2 molecular diagnostic used to detect viral RNA in clinical samples *(2)*. This assay is widely used and also serves as a prototype for many FDA-authorized tests (discussed below). As I show here, this assay uses a poorly-designed specimen extraction control that does not perform as intended on real-world samples. The CDC control fails to reports on specimen and process integrity.

I alerted the CDC to this potential problem in March of 2020, supported by bioinformatic prediction. Their response, sent more than a month later, did not share my concerns. One, the CDC had not received widespread reports of false-negative tests. Two, negative patients should err on the side of caution and re-test if concerned. Three, the FDA had approved this test, making it, *de jure* correct (full correspondence in Supplemental Materials).

False negatives are inherently difficult to detect – particularly when the assay in question is the current gold standard. Testing reagents are in short supply, making it hard for individuals to be tested once, much less repeatedly. Official approval is not a defense against facts. As I show below, the CDC design is both flawed and fixable. Individuals would be incensed by a preventable false-positive diagnosis putting them into unnecessary quarantine. The public should be incensed by a poorly designed official assay whose preventable false-negative diagnoses allow virus-positive patients back into society.

## Results

The 2019-nCoV Real-Time RT-PCR Diagnostic Panel is a CDC-designed and validated COVID-19 assay comprised of three qPCR primer/probe sets. N1 and N2 generate SARS-CoV-2-specific amplicons from reverse transcribed viral RNA. The third, RP, targets the human *RPP30* gene and is an “extraction control” that is intended and claimed to inform about successful sample collection, RNA extraction, and reverse transcription *(3)*. It does not perform these functions in practice. The CDC design is fundamentally flawed and has the potential to return dangerous false negatives (Fig. 1A).

**Fig. 1.**
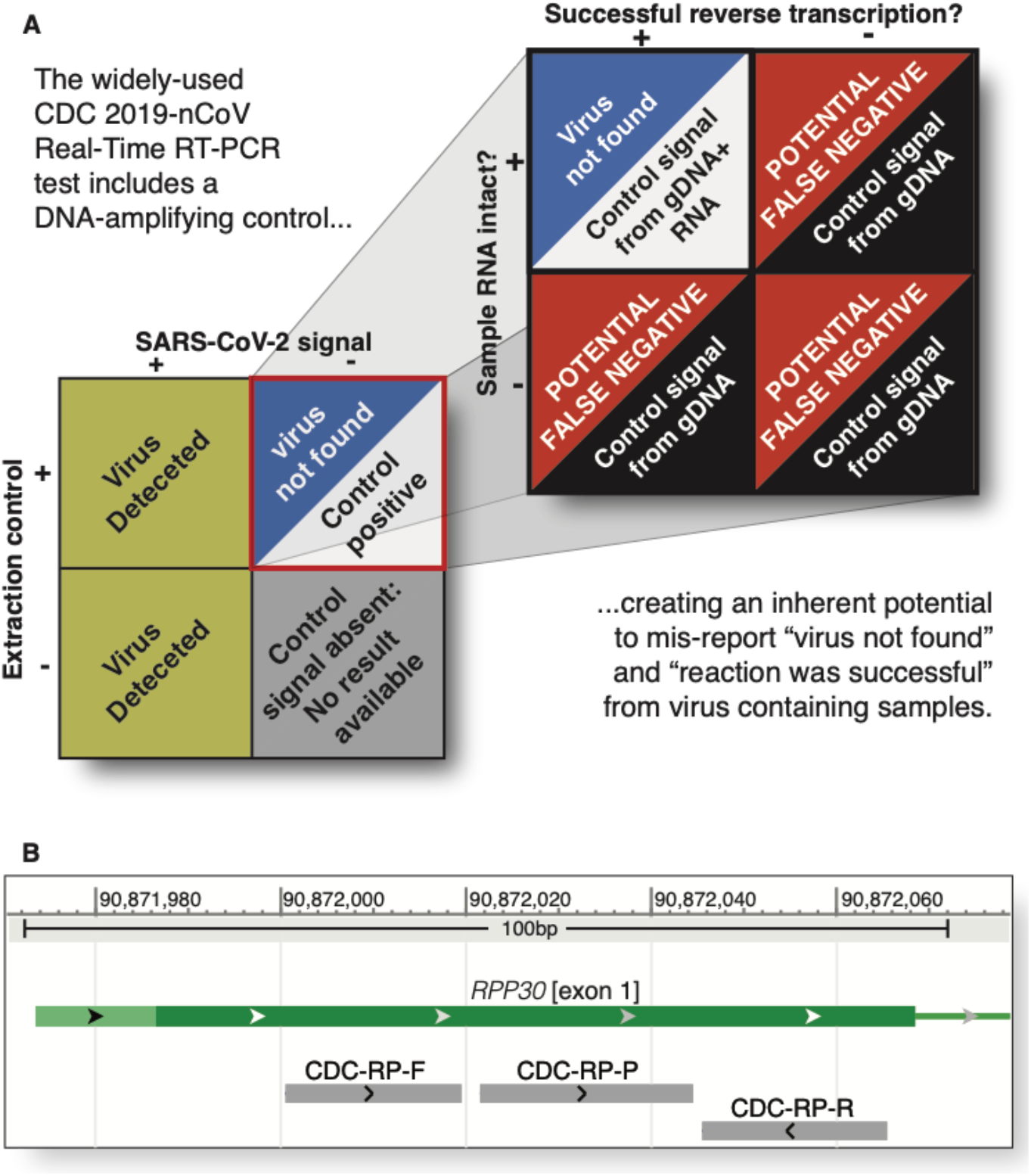
**A**. The CDC-specified SARS-CoV-2 qPCR assay and associated clinical algorithm are intended to inform for presence of virus, or that the assay worked and could have found virus, but that no virus was found. Signal from virus-targeting RT-qPCRs is sufficient to report “Positive 2019-nCoV”. An “invalid” report is returned where all viral and control probes are negative. The clinical interpretation is “2019-nCoV not detected” when a human control, targeting the *RPP30 gene*, returns a positive signal and when viral probes are negative. Patients who are truly virus negative will fall into this category. So will patients for whom sample collection, handling, or analysis problems have occurred, leading to false negative results. **B.**The CDC-specified extraction control primers and probe recognize a sequence that is identical in reverse transcribed RNA and genomic DNA. Megablast search of the human refseq_dna database identifies perfect matches for the CDC-control primers, RP-F and RP-R, and probe, RP-P. This design amplifies and detects a 65 base pair amplicon located entirely within exon 1 of the *RPP30* gene, located on human chromosome 10 (coordinates from *hg38* human genome build). Specific detection of reverse-transcribed RNAs can be accomplished by design of an exon-exon junction spanning amplicon and probe (Supplemental Figure 3).

The CDC-specified RP extraction control design generates identical amplicons from reverse transcribed RNA and human genomic DNA. Megablast *(4)* search of the RP forward, reverse, and probe sequences against the human refseq_rna transcript database identifies perfect matches within the *RPP30* mRNA. The design recognizes sequences that are entirely contained within a single exon, and therefore generates an identical amplicon from the *RPP30* genomic locus (Fig. 1B). The CDC extraction control generates a positive signal in reactions containing intact RNA. It also generates a positive signal from RNA-free reactions containing human genomic DNA (Fig. S1).

Genomic DNA is co-purified in quantities sufficient to generate strong positive signals for the CDC-specified extraction control during work-up of clinical RNA specimens. To test for the presence of control-affecting DNA, qPCR reactions lacking reverse transcriptase were performed on SARS-CoV-2-positive clinical samples using the CDC-specified RP primer and probe. “Purified RNA” samples from COVID-19 cases were obtained from a CAP/CLIA certified university hospital molecular pathology laboratory. Samples had been processed using either solid phase extraction (Qiagen) or low pH organic extraction (Trizol). Both methods are permitted under current Emergency Use Authorization (EUA) for RNA isolation. In reality, both methods co-purify significant amounts of DNA *(5)*. All clinical samples tested generated unambiguous extraction control positive signals in the absence of reverse transcription, a reaction context that could not have detected virus an RNA virus (Fig. 2A)

**Fig. 2.**
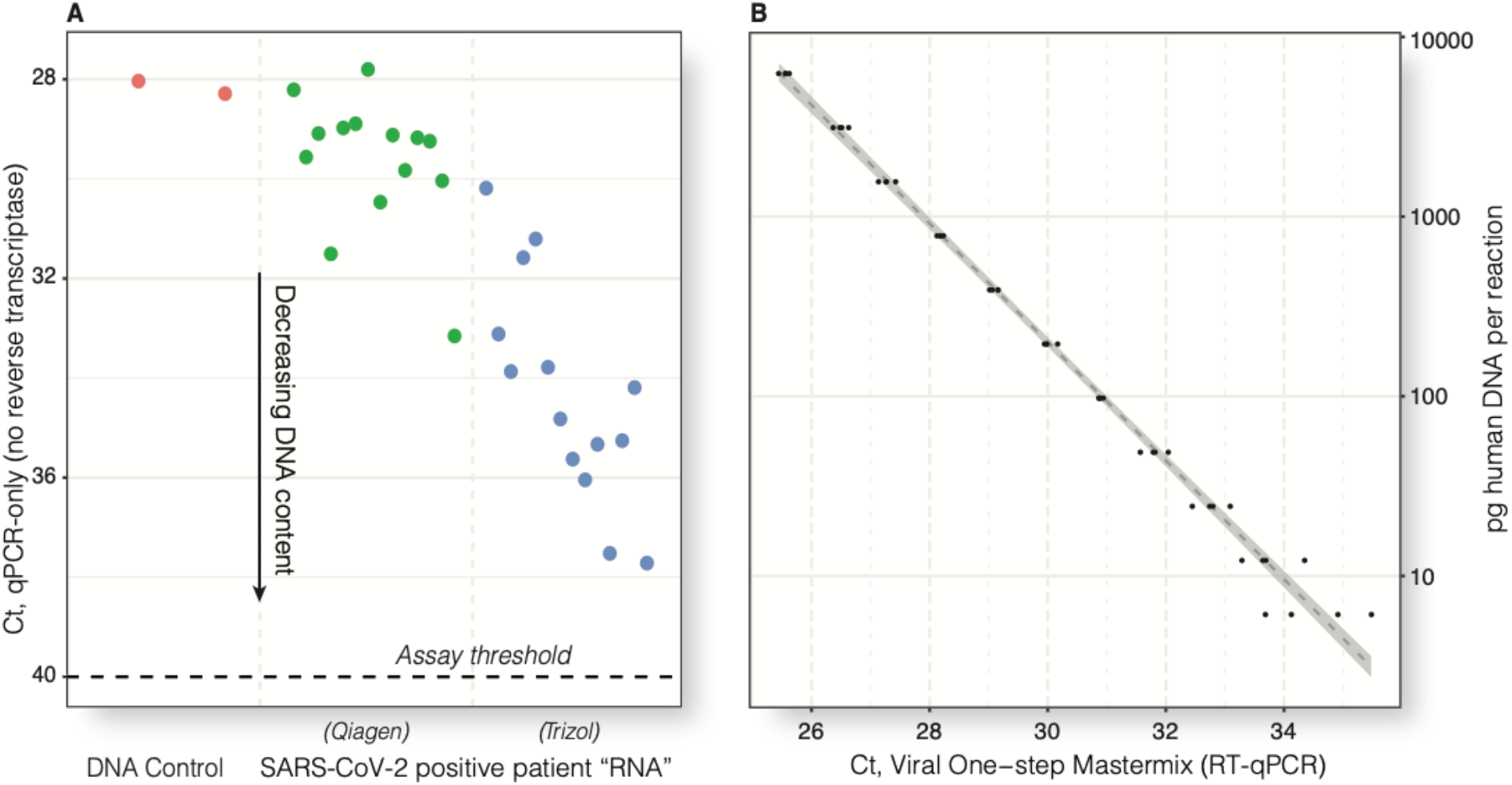
Genomic DNA is co-purified during RNA extraction of clinical COVID-19 samples and is sufficient to generate a positive signal using the CDC extraction control. **A.**qPCR (without reverse transcription) was performed using the CDC extraction control (RP) probe. Raw Ct values are shown. A total of 28 representative COVID-19 positive clinical samples were tested, fourteen purified by automated silica spin-column (Qiagen) and fourteen by manual organic extractions (Trizol, Materials and Methods). All samples contained genomic DNA and generated extraction control positive signals well above the CDC assay threshold (40 cycles, horizontal dashed line). Significantly more genomic DNA was present in samples purified by silica spin-columns compared to organic extraction (mean Ct of 29.6 and 34.3, respectively, *p*= 1.21×10^−6^, unpaired *t*). Co-purification of DNA is to be expected based on the mechanisms of purification used, and is openly caveated in vendor literature *(13)*. Per-plate process controls, shown, contained 8.6 ng of genomic DNA, equivalent to ~1250 diploid cells. **B.**Single-cell equivalents of purified genomic DNA, absent RNA, are sufficient to generate an “extraction control positive” signal using the CDC-RP probe. RT-qPCR was performed using Viral One-Step Mastermix (Materials and Methods) on 2-fold serial dilutions of a commercially-supplied reference human DNA (RNA-free). Dilution equivalent to one diploid copy of gDNA per reaction (6.125pg) generates signals well above the CDC-specified threshold, despite increase in Ct variability at extreme dilutions.

Single-digit copies of genomic DNA are sufficient to generate a positive control signal using the CDC-designed assay. Driven by concern about the control probe signals observed in PCR-only analysis, I determined the quantity of genomic DNA required to generate an assay-positive control signal. Commercially supplied, pooled human DNA was serially diluted across several orders of magnitude and analyzed by the same one-step RT-qPCR used for clinical samples (Fig. 2B). Extrapolating from pure-DNA samples, a positive signal from the CDC control would be readily achieved in samples containing one copy of the genome copy per reaction *(7)*.

Due to the presence of co-purifying genomic DNA in clinical samples, loss of RNA integrity leads to false-negative results using the CDC-specified control. RNA, including the genome of the SARS-CoV-2 virus, is intrinsically more labile than DNA. RNA-degrading enzymes (RNases) rapidly hydrolyze RNA while leaving DNA intact. Laboratory procedures that manipulate RNA, including methods specified in the CDC EUA, require careful sample, reagent, and consumable handling to avoid introduction of ubiquitous environmental or personnel-derived RNases. Loss of RNA integrity, accomplished here by intentional exposure to RNase, causes virus-positive clinical samples to return a false-negative “extraction control positive, virus negative” result using the CDC-design (Fig. 3).

**Fig. 3.**
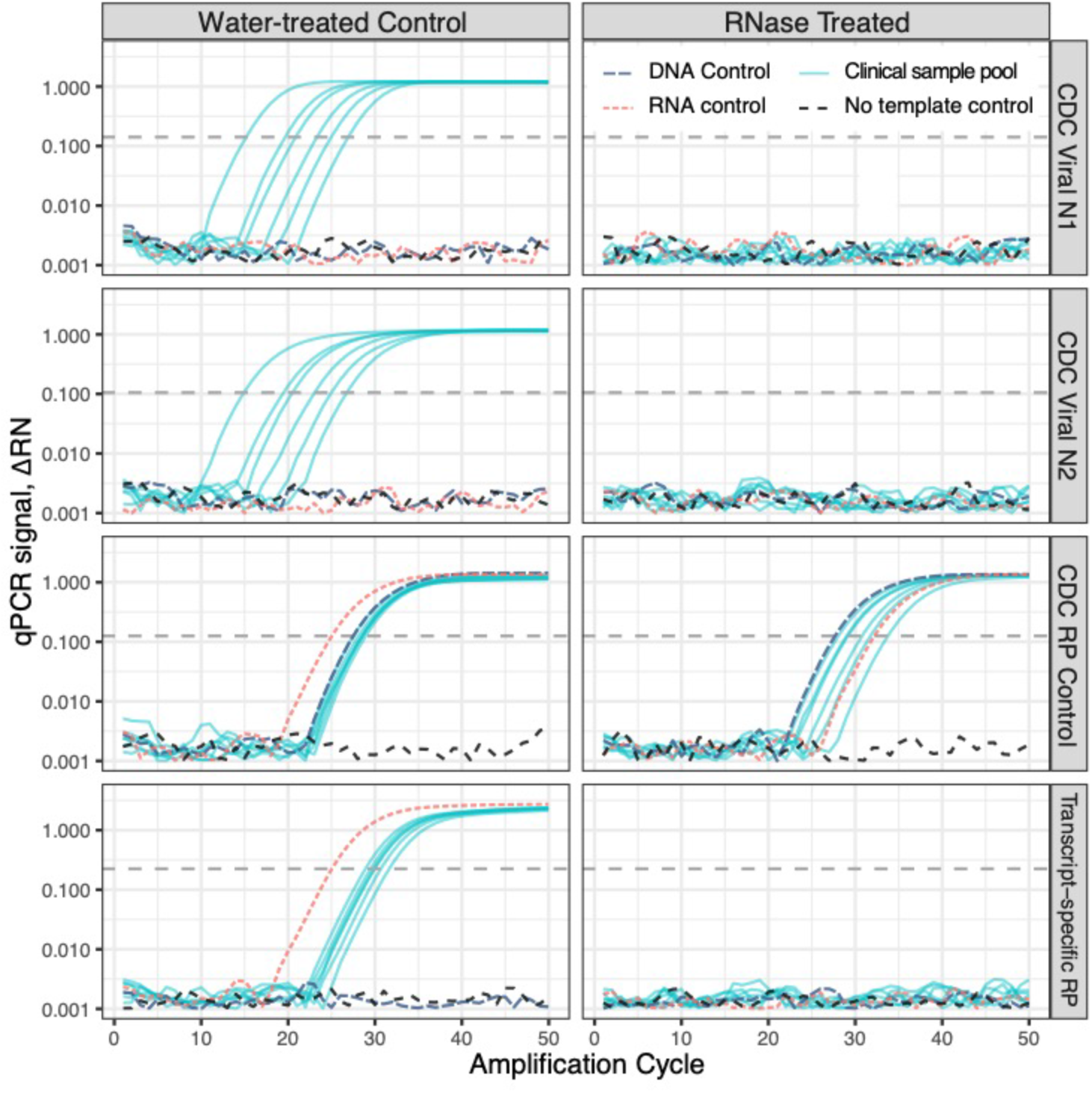
CDC-specified testing of SARS-CoV-2-positive clinical specimens return a false negative following loss of RNA integrity. Reverse-transcriptase-qPCR data were generated using Viral One-step RT-qPCR mastermix (Materials and Methods). Extracted nucleic acid samples used in Figure 2 were pooled, divided into aliquots, and treated with either nuclease-free water or Ribonuclease A (RNase A). Commercial RNA, DNA, and no-template controls (NTC) were treated in parallel. Samples were analyzed using the Viral N1, Viral N2, and CDC-designed RP control primer/probe sets, and a transcript-specific control targeting exons 1-3 of *RPP30 (Table S1)*. Correct positive viral and control signals are generated from each infected-patient pool after mock treatment (water, left). Loss of RNA integrity causes clinical samples to return negative signals from both viral probes, but the CDC-specified RP control due to DNA-templated amplification (RNase, right). A transcript-specific RP control accurately informs as to loss of sample integrity. As desired, samples with degraded RNA do not generate an extraction control signal from a transcript-specific design, marking the result as invalid, rather than “virus not detected”. Horizontal dashed lines reflect software-generated Ct thresholds.

Assays can be designed to specifically detect mature mRNA, and ignore genomic DNA, by generating amplicons that are of PCR-amplifiable size only after splicing or by targeting sequences created at exon-exon junctions (Figs. 3, S1-3). A transcript-specific control design incorporating both of these approaches correctly monitors human specimen collection, RNA integrity, and successful reverse transcription, thus serving all intended roles of the human specimen / extraction control (Figs. 3, S1, and S2 and Supplementary Text)

## Discussion

Properly-designed extraction and reverse transcription controls are essential, not for detecting SARS-CoV-2, but for determining when a negative test result is invalid. The unprecedented scale of molecular testing undertaken during the COVID-19 pandemic has required that labs rapidly adopt this CDC-specified, among many other, new assays. Labs accustomed to automated walk-up systems are now performing manual sample handling and reaction setup one pipet tip at a time. The CDC design detects SARS-CoV-2, but in the face of real-world assay errors, its faulty control design fails-unsafe and calls into question virus-negative results.

The CDC specifies a once-per-plate control using *in vitro* transcribed RNA, purified virus, or pooled patient samples. This approach does not inform about the suitability of individual specimens. Degradation of sample RNAs or failure of reverse transcription on a per-sample basis results in the inability to detect viral RNAs while the DNA-templated control signal remains. This opens the door to false-negative clinical interpretation of patients from whom virus was successfully collected.

Controls for specimen integrity and success of all reaction steps are critical to robust assay design. The currently-specified CDC control achieves neither. A number of other FDA-approved SARS-CoV-2 molecular assays use synthetic RNA spike-ins added at the time of extraction. This approach controls for post-extraction sample integrity and reverse transcription, but does not inform about sufficiency of collected patient samples *(9)*. The intent of the CDC’s extraction control is good, monitoring (1) sufficient human specimen collection, (2) maintenance of sample integrity during processing, and (3) successful reverse transcription and qPCR. These goals can be met by an extraction control that specifically detects human RNA and is insensitive to genomic DNA; the CDC’s mis-executed plan can be put into practice by manufacturing and shipping an updated control.

The DNA cross-reactive control probe is used in over 40% of the COVID-19 FDA-authorized protocols listed as of May 13, 2020 *(10)*. 31 million assays worth of CDC-design reagents have been produced and shipped by a single vendor as of April 22, 2020 *(11)*. This CDC-designed SARS-CoV-2 assay is FDA-approved and unquestionably popular, but we must not confuse popularity and official approval with being correct. The scope of this problem is large, though its extent is unknowable: the sequences used in many test designs, commercial and academic, are not disclosed to the public. Even fully disclosed sequences do not guarantee rapid discovery of errors. This poorly-designed CDC extraction control probe has been included in clinical RT-qPCR panels used in testing for RNA viruses across more than a decade *(12)*. Past errors cannot be unmade, but future false negatives that would arise from this design can be prevented.

The current CDC algorithm states that molecular testing “…must be combined with clinical observations, patient history, and epidemiological information”*(2)*. In the context of drive-up testing and potential nationwide return-to-work testing, clinical observations will be necessarily limited and often impersonal in scope. Absence of data and data of absence must be carefully distinguished. A proper control does not provide definitive diagnostic results from a flawed specimen. It will, however, fail-safe, flag a result as invalid, and call attention to problem samples to enable focused re-testing or changes in clinical management.

In the interest of public health, assays must generate as many valid results as feasible. This requires simultaneously maximizing identification of true positives *and* minimizing false negatives. Virus-shedding patients must not be given an all-clear test result due to an improperly designed that can be readily fixed. While perfect diagnostic tests are unachievable, assays should be designed and updated to strive for maximum specificity.

The cost of incorrectly clearing SARS-CoV-2-positive patients is incalculably large. Implementing a redesigned control that performs as intended amounts to pennies per test. A new primer / probe set, such as the one described here, is a drop-in replacement that can be manufactured rapidly and at scale. It is imperative that the CDC take action to update its clinical algorithms and specify a re-designed transcript-specific extraction control that performs all of its claimed, and critically important, functions.

## Acknowledgments

APR would like to thank Amy A. Caudy for technical discussion and critical reading of the manuscript, Bruce Futcher for discussion, and Richard Kew and Karen Bai for assistance accessing clinical samples. APR would like to acknowledge the support of the Stony Brook Medicine Biobank for providing de-identified purified RNA from patient samples and Maple Flavored Solutions LLC for access to instruments and donation of supplies and reagents.

**Funding** the author is supported by startup funds from Stony Brook University and NIGMS R01GM132238

## Author contributions

APR conceived, designed, executed, analyzed, and wrote this manuscript

**Competing interests** the author declares no competing interests

**Data and materials availability** all data are available in the manuscript or the supplementary materials.

## Supplementary Materials

Materials and Methods

Supplementary Discussion

Figures S1-S3

Table S1

## Materials and Methods

### Sample preparation

Fully de-identified clinical “RNA” (more properly, total nucleic acid) specimens, generated in excess during testing of SARS-CoV-2 RT-qPCR-positive patients, were provided by the Stony Brook Hospital Biobank. Nucleic acids had been previously purified from polyester-tipped nasopharyngeal swabs desorbed in viral transport media (VTM) and processed in a CLIA high complexity pathology laboratory. Briefly, nucleic acid was extracted from 160μl VTM using Qiagen QiaAMP DSP Viral Mini spin columns processed on a Qiacube automation system as per vendor instructions (Qiagen, BV) or from 200μl VTM using manual Trizol-mediated organic extraction (Life Technologies) followed by alcohol precipitation *(5)*. Samples were eluted in 50μl of RNase-free water supplemented with 0.04% NaN3 (Qiagen) or resuspended in RNase-free water (Trizol) and stored at −80C until use. Commercial pooled RNA (Quantigene Human Reference, QS0639, Life Technologies) and pooled human male DNA (G3041, Promega and 4312660, Life Technologies) were used to generate calibration curves and serve as reaction controls. DNA samples were diluted in nuclease free water supplemented with 0.01% Tween-20, RNA samples were diluted in nuclease-free water. Low binding filter tips were used for all liquid handling.

### PCR primer and probe design

qPCR primers and hydrolysis (“TaqMan”) probes were used for real-time detection. CDC-specified RnaseP (RP) and SARS-CoV-2 (N1 and N2) probes were purchased from Integrated DNA Technologies (RUO qualified, 10006713; to conserve clinically certified reagents, research grade primers of identical composition were used). CDC primers were used as specified in the EUA

A total of five *RPP30* probe set designs from multiple vendors were ordered and empirically tested (Table S1). These include probes targeting exon-exon junctions and a minor-groove-binding TaqMan probe (Life Technologies). The best performing design, generating an amplicon across exons 1-3 with an exon-exon junction-spanning probe, was ordered with a 5’ 6-carboxyfluoroscein, 3-BHQ and internally-quenched probe from IDT as Assay Hs.PT.58.2854947. This design had the highest sensitivity (as lowest Ct), highest efficiency, and best consistent terminal signal characteristics of all candidate probes tested (Supplementary Figure S1a). All non-CDC primer/probes were used at 10 pmole primer/5 pmole probe per 20μl reaction.

### qPCR reaction conditions

Reverse transcription coupled qPCR was carried out essentially as described in the CDC EUA for COVID-19 testing *(2)* with the following modifications. To conserve supplies for clinical use, comparable research-grade reagents were used where possible. TaqMan™ Fast Virus 1-Step Master Mix (4444432, Life Technologies) was used for one-step RT-qPCR and TaqMan Fast Advanced Master Mix (4444556, Life Technologies) was used for non-reverse transcriptase reactions. Real-time data were collected on a validated ViiA7 qPCR instrument using a 384w block and vendor plates and seals. All runs were performed using CDC-specified thermal cycling parameters as per the EUA. Automatic baselining and thresholding of raw qPCR data was performed using vendor software (QuantStudio 1.3, Life Technologies).

### Measurement of qPCR efficiency

Efficiency of all qPCR probes was tested by using a simplex 2-fold serial dilution of Quantigene commercial reference human RNA (Invitrogen) or DNA (Promega) ranging from 64.2ng to 62.7pg per reaction using Virus One-step Mastermix and Taqman Fast Mastermix. To robustly determine CDC-RP response to genomic DNA, four completely independent dilutions of genomic DNA were performed down to 7.8pg/reaction and analyzed using Virus One-Step Mastermix.

### Sample pooling and RNAse treatment

Limited-volume clinical samples were pooled to generate sufficient material for the RNase-exposure experiment. A total of 28 clinical RNA samples were pooled, fourteen each purified by Qiagen and Trizol (above, <u>Sample preparation</u>). Three pools were generated for each technology, containing 5, 5, and 4 clinical samples, respectively. Pooling was performed based on sequence of the de-identified serialized sample names (Q1, Q2…Q14) provided by the Stony Brook Hospital Biobank. Sample names do not reflect time of collection nor any other patient or sample-intrinsic information. Pool composition, beyond positive SARS-CoV-2 status and by extraction technology, is effectively random.

8μl of each clinical purified nucleic acid specimen was pooled into a fresh RNase-free polypropylene tube. 8μl of nuclease-free water was added to each of the four-specimen containing pools to compensate for the lower specimen count (resulting in a total volume of 40μl per pool). 20μl of each pool was aliquoted to a fresh tube. 20μl aliquots of 1.73ng/μl Promega control DNA and Quantigene RNA were generated, as were 20μl aliquots of nuclease-free water as a no-template control. 1μl of nuclease free water was added to each of the “sham” samples (six pools plus three controls). 1μl (0.7U/μl) of recombinant RNaseA was added to each “RNase” sample in a physically separate laboratory space. Capped tubes were mixed, vortexed, centrifuged briefly, and incubated at 37° for 90 minutes immediately prior to RT-qPCR setup. One-step RT-qPCR was performed on treated and control samples using CDC-RP, CDC-N1, CDC-N2, and Hs.PT.58.2854947 assays.

### Plotting of raw qPCR data

Raw qPCR data were exported as deltaRN (ROX-normalized, background subtracted fluorescence) per cycle using QuantStudio. “AmplificationData” were imported into R for plotting. Optical and quantization noise result in small values centered around 0 (i.e. either + or − 1×10^−4^) during cycles prior to generation of detectable signal. To eliminate undefined log10 transformation errors of negative values, a small offset value of 0.004 was added to each DeltaRN signal prior to plotting on log-axis scale. This is evident as a displayed baseline signal centered around 0.004 in Figure 4. Unmodified data were used for all calculations and are included in Supplementary Data; this transformation is for convenience to make ≤0 data plottable on a logarithmic axis.

## Supplementary Text

### Controls are useful; why not add more?

Assay design should be fit to task and balance throughput, cost, and sensitivity/specificity. Controls are necessary, but add assay complexity, reagent cost, and can compete with throughput. The current CDC-EUA uses a single detection channel, 6-carboxyfluoroscein, for each of three probes that are analyzed separately. Three wells are required per patient or control specimen. Adding even one more control decreases assay capacity by 25% while increasing the cost of One-step mastermix per patient by the same amount. More is not always necessary. A well designed, transcript-specific sample collection, extraction, and RT-qPCR probe can achieve a composite control using one channel. If the control does not turn positive, a virus-negative reaction cannot be trusted. A composite control does not specify where a problem occurred, only that one has.

Modern qPCR (and by extension, RT-qPCR) is not limited to a single detection channel; multiplex designs can generate data from two to five (or more) targets in a single well. Multiple fluorescent dyes on probes of different sequence can be combined in carefully-designed assays, increasing the number of targets that can be interrogated in one tube. Limitations of multiplex qPCR hardware, fluorescent dyes, and the realities of molecular biology constrain the degree of multiplexing and drive design concessions. Multiplex designs are widely used in the clinic, including for SARS-CoV-2 testing. The ability to multiplex does not obviate good assay design. Several current panel designs use a synthetic RNA spike-in to monitor extraction and reverse transcription/PCR. Patient sampling and extraction controls are critical to clinical interpretation and should be prioritized. As of May 12, 2020, none of the FDA EUA’d SARS-CoV-2 assay designs contain *both* a human sample collection control and a separate extraction/RT control.

Four-plex RT-qPCR, used by several panels, could be leveraged as a pair of viral probes, an RNA spike to monitor extraction/RT-qPCR, and an RNA-specific human specimen control. This approach retains control for presence of a human specimen while providing more granularity than available from a single composite control, such as a transcript-specific *RPP30* probe. This two-control design would enable more efficient handling of specimens where no human specimen control signal is generated. An intact spike-in signal would mean that that extraction and downstream handling of RNA through RT-qPCR were successful, despite no patient RNA having been obtained. Patients should be called back in, re-sampled, re-extracted, and re-tested. In contrast, failure of the spike-in control means that something likely went amiss during handling. Whether caused by a bad pipet tip, a one-off problem with extraction, or shaky hands at the end of a shift, excess viral transport media could be re-extracted and re-tested before recalling a patient into the health care setting for a re-swabbing.

### Why change the assay when one could fix the input?

Although a re-designed transcript-specific control eliminates the false negative potential described here, approaches to make best use of the current reagent will likely be proposed. One potential alternative to eliminate confounding genomic DNA is to digest extracted samples with deoxyribonuclease (DNase). This should be avoided. This approach is expensive (far more than a re-designed primer), requires addition of divalent cations that can interfere with downstream reverse transcription and PCR, and is imperfect: DNA has evolved to be more physically robust than RNA. As shown above and in many other contexts, exponential amplification during PCR generates detectable signals from single-digit copies of template. Even commercial offerings “reference RNA” can be contaminated with low, but interfering, levels of human DNA. Quantigene human reference RNA (Life Technologies) is specified as a mixture of DNase-treated RNAs from multiple cell lines. In practice, this material continues to generate qPCR signals both using a Taq-only, non-reverse transcribing, mastermix and a one-step RT-PCR mix following RNase A treatment (Figs. 3 and S2).

### Ct is a unitless measurement and should be used with care when discussing quantity

qPCR is able to operate in qualitative, semi-quantitative, or fully quantitative modes, depending on assay design. Currently deployed assays are effectively qualitative (Yes/no. Above or below acceptable Ct). Throughout this manuscript, absolute Ct values and difference in Ct between samples are used as a proxy of inverse(log2) amount of input material or differences thereof between samples. Comparisons are restricted to same-probe, same-instrument, same (or similar) reagent. Calibration of Ct:[nucleic acid] was performed using dilutions of commercial standards. An exception to this conservative approach is the use of the “40 cycle” threshold specified in the CDC EUA. Our results err on the side of underestimating the extreme sensitivity of the primary CDC design. The Applied Biosystems Viia7 384w instrument used here is moderately *less* sensitive (higher Ct) for a given reaction composition and volume than a 96w FAST reaction block in the same instrument (information obtained from Life Technologies technical support, 2020/05). For a representative instrument, the 96w FAST block Ct was approximately 0.6 cycles lower across the dilution range.

### Assays should be improved when feasible, not rubber-stamped and set in stone

I alerted CDC that their EUA-specified suffered from a potential false-negative on March 20, 2020 (attached below). Their response, sent thirty-six days later (also attached), dismissed my concerns at multiple levels. One: there had not been widespread reporting of false negatives from this test. Two: CDC’s clinical algorithms state that negative-testing patients should be re-tested and that clinical management should not rely solely on this test. Three: the FDA had approved the test, making it, *de jure*, acceptable.

Each of the CDC’s arguments is fallacious. False negatives are inherently difficult to catch, and identification of true positives is a challenging bootstrap problem. The idea of “re-test if you’re uncertain” is tone deaf in the context of widespread shortages of testing reagents and long backlogs for sample processing, not to mention the moderately invasive nature of nasopharyngeal sampling. Appealing to FDA’s authority may be legally satisfying, but does nothing to actually address issues of scientific merit.

We will never know how many virus-positive patients have been given what amounts to an “all clear” due to false-negatives generated by this flawed control design. The potential for misdiagnosis is real, and there is a window to make an important assay work as it was intended. The alternative would be for CDC to say “our test doesn’t work as we intended; if you are worried that you’re sick and your test has come back negative, come back and be tested again until your fears are allayed, or you test positive.”

Fixing some problems is difficult; others are difficult to identify but easy to remedy once found. As shown here, this important assay can be made to operate as intended by a re-manufacture of a single reagent.

---

**Fig. S1.**
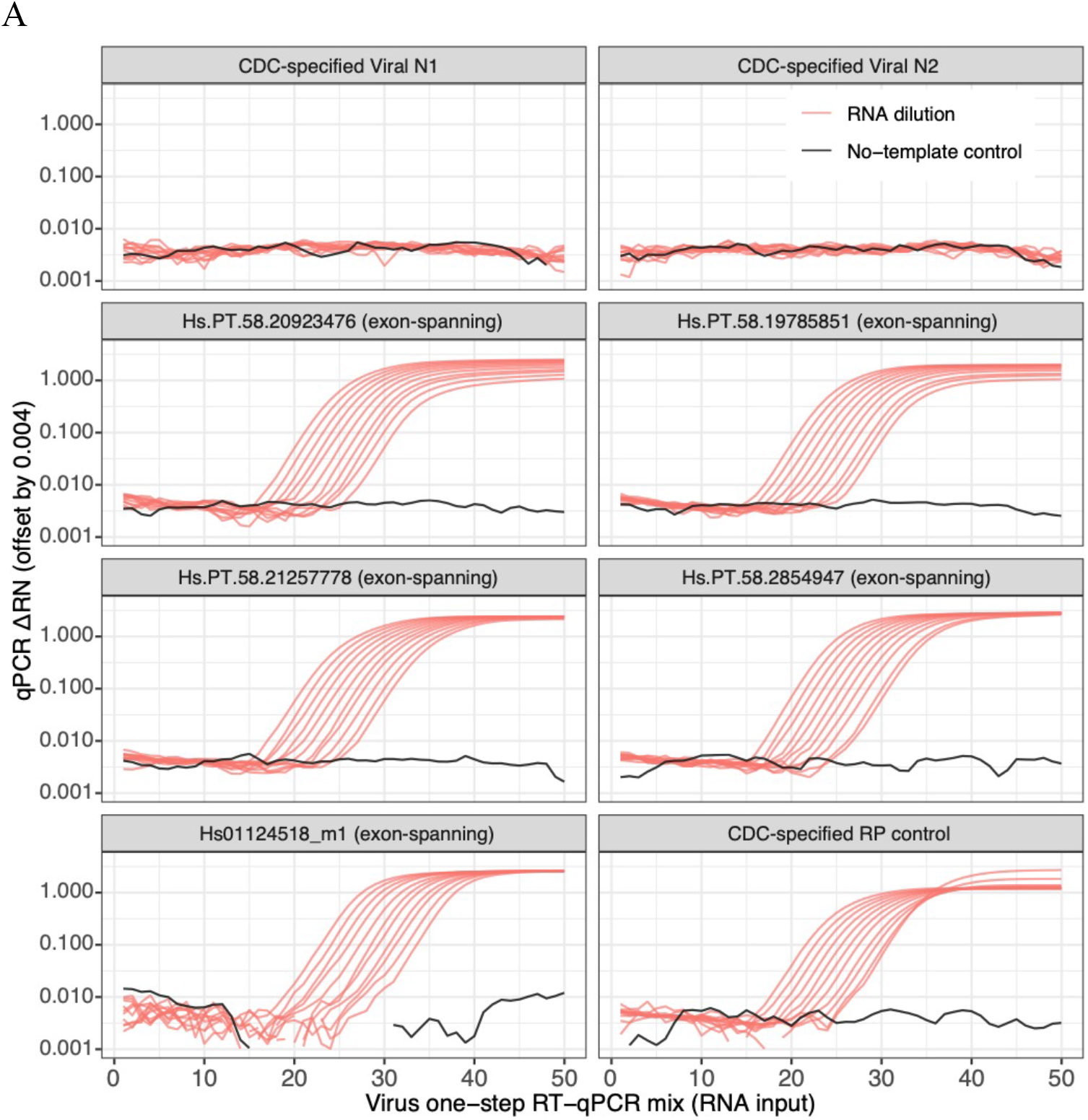
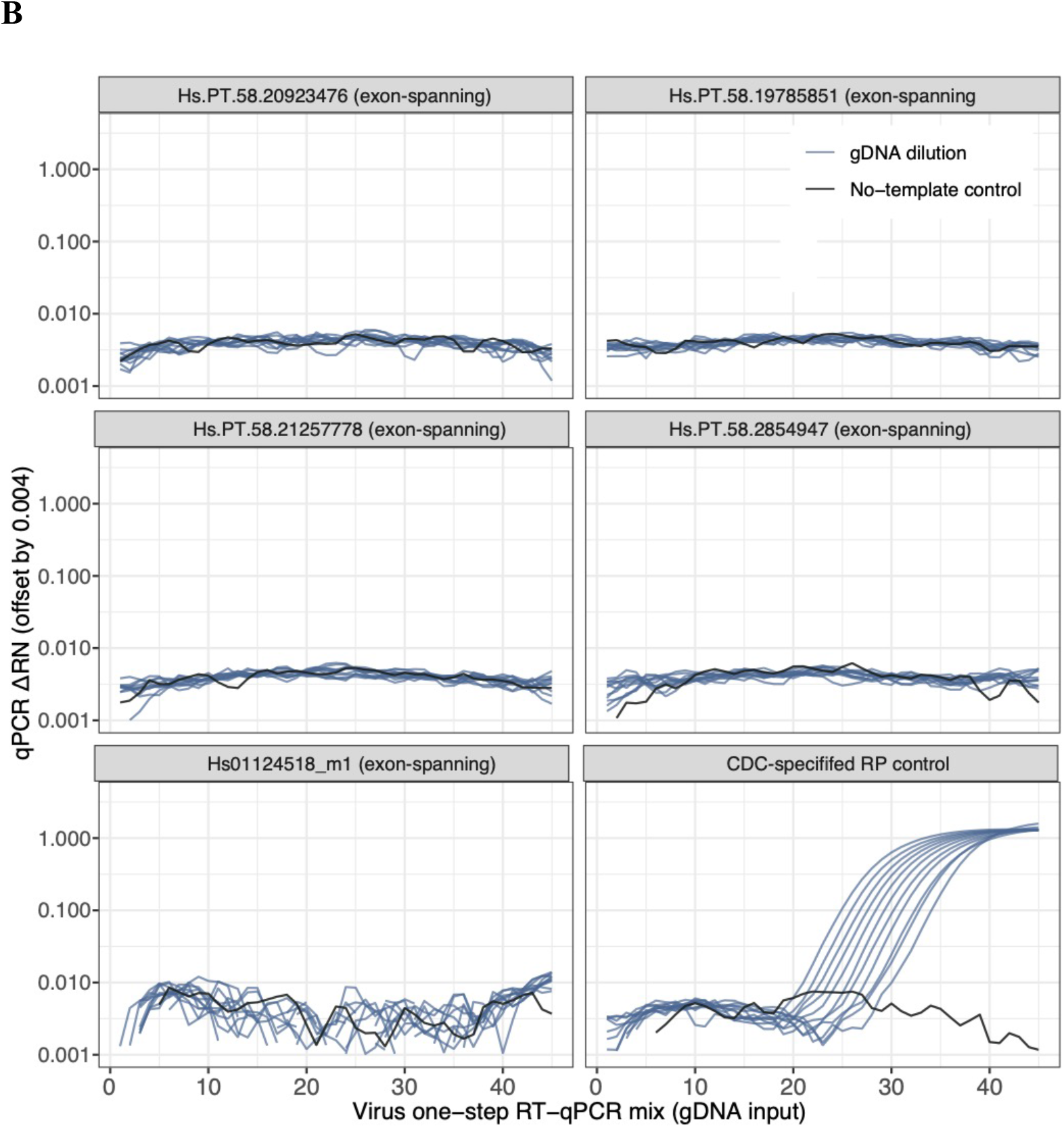
RNA, but not genomic DNA, is a productive template for reverse-transcription-qPCR amplification using any of the five tested transcript-specific *RPP30* controls. One-step RT-qPCR was performed with the same mastermix used for clinical samples across a 2-fold dilution series (Materials and Methods). As expected, pre-pandemic commercial reference RNA is non-reactive to the SARS-CoV-2 primer/probe sets. All six primer/probe sets targeting *RPP30* show a strong and relatively linear signal across a range of RNA input (A) (S1a). Using the same reverse-transcriptase containing mastermix with mass-equivalent dilutions of RNA-free genomic DNA input, none of the transcript-specific probes generated a qPCR signal, within 50 cycles, across a titration of purified DNA ranging from 20,000 copies to 19.5 copies/reaction. **(B)**The CDC-specified RP control generates a strong signal from genomic DNA. Reactions containing 64ng of either total RNA or genomic DNA generated a ~3 cycle difference for threshold crossing (Supplemental Text). Modest baseline and NTC signal is due to offsetting of raw qPCR signals before plotting, as described in Materials and Methods.

**Fig. S2.**
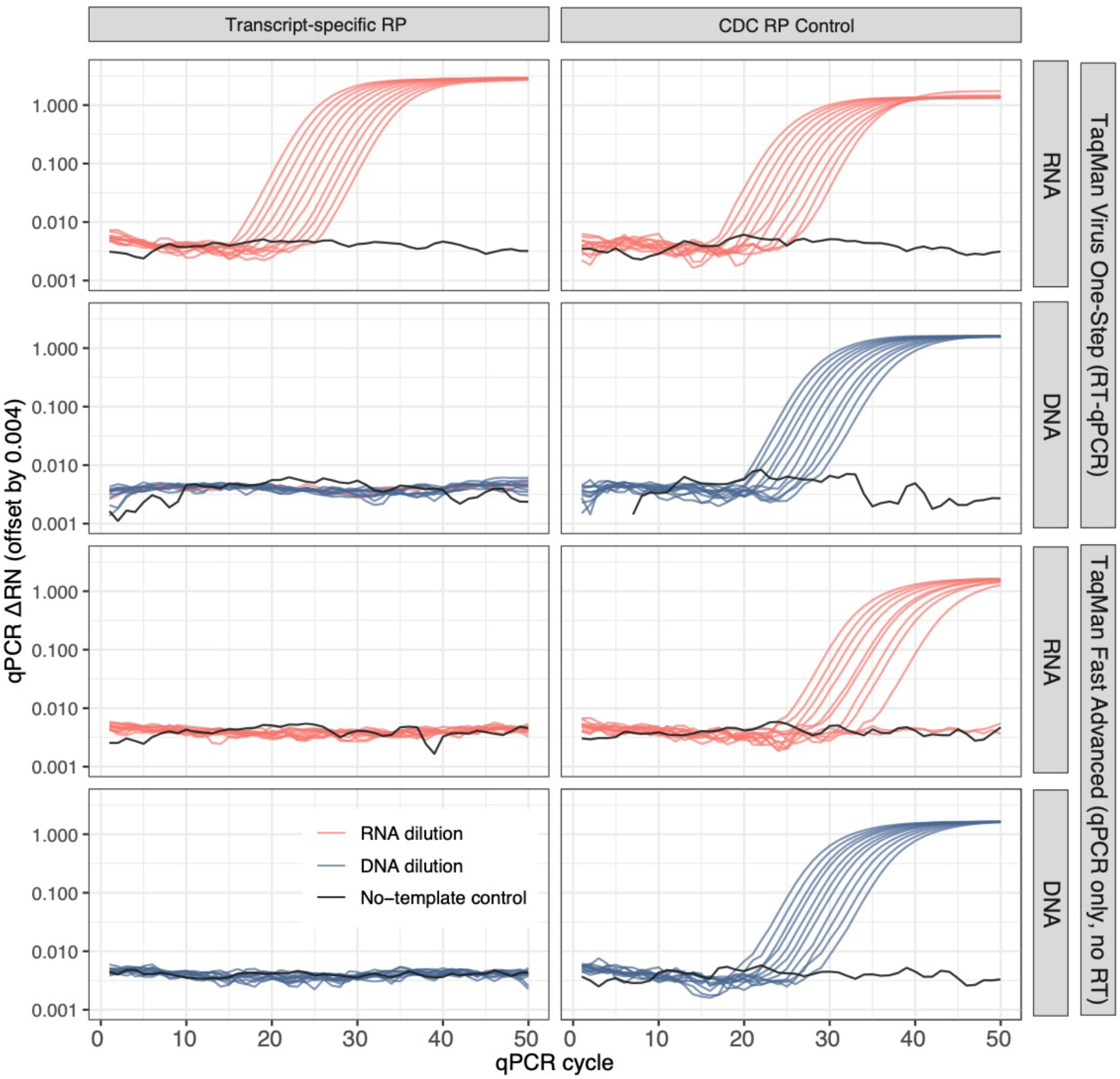
Positive signal from a transcript-specific *RPP30* probe requires both RNA and reverse transcriptase (RT). The CDC-specified RP control requires neither RNA nor RT when genomic is present. Dilutions of either commercial RNA or DNA were used as input for reactions containing either Viral One-step RT-qPCR mastermix (containing reverse transcriptase) or TaqMan Fast qPCR mastermix (lacking reverse transcriptase) using either the CDC-specified or transcript-specific *RPP30* probe. The transcript-specific probe generates signal only in reactions with an RNA input where reverse-transcriptase is present (as a one-step mastermix). Genomic DNA does not generate a signal above baseline even in an extended 50-cycle amplification, nor does RNA generate a signal in a qPCR-only (no reverse transcriptase) reaction for a transcript-specific probe. Signal generated for CDC-RP probe in PCR-only analysis of “RNA” is due to low-concentration genomic DNA present in Quantigene commercial RNA (supplemental discussion), further underscoring the need for transcript-specific primer design.

**Fig. S3.**
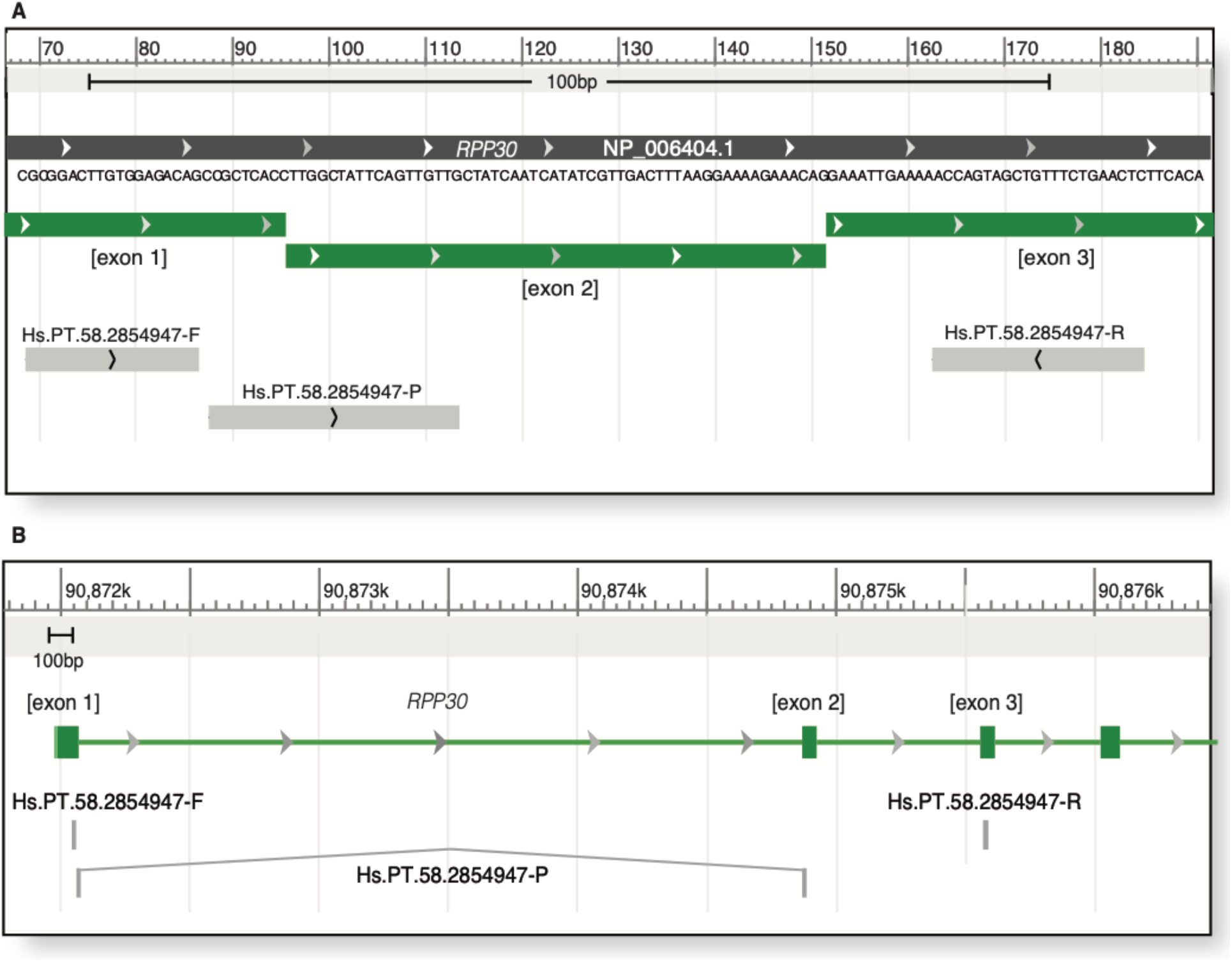
The cDNA-specific *RPP30* assay used here is doubly-unable to generate signal qPCR signals from genomic-DNA. Reverse-transcribed *RPP30* RNA can serve as a template for the Hs.PT.58.2854947 qPCR assay generating a relatively short amplicon (S3a). The forward and reverse primers bind in the bodies of exon 1 and 3, respectively. The hydrolysis probe binds at the exon 1 – exon 3 junction that is formed during mRNA splicing. This primer/probe set does not generate a signal from a genomic DNA template (shown above), due to the ~3.5kb spacing between the forward and reverse primers that is un-amplifiable using the short annealing/extension cycles of the CDC ramping program (S3b). Even if PCR product were generated, the hydrolysis probe binding sequence is split in half, with each side ~3kb apart in a genome-templated amplicon.

**Table S1.**
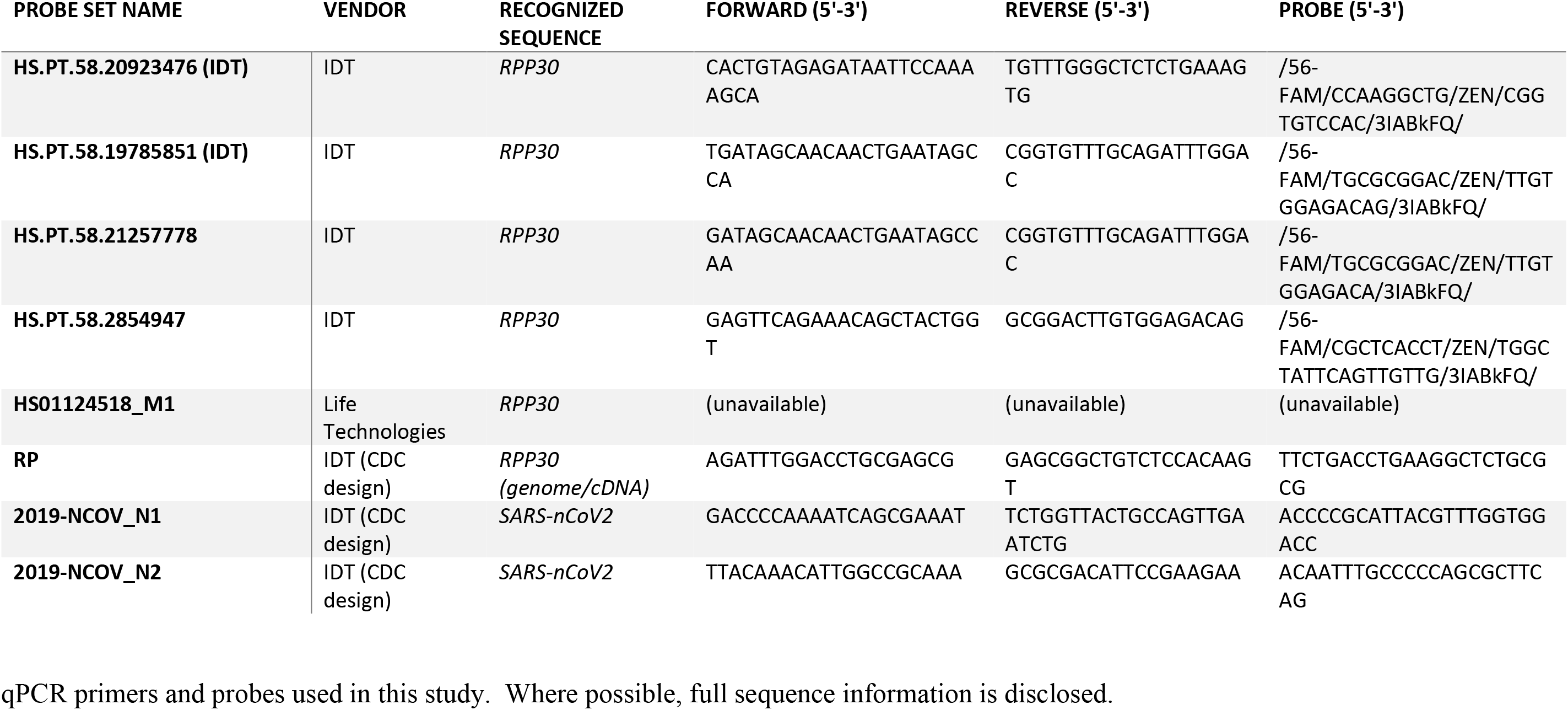
qPCR primers and probes used in this study. Where possible, full sequence information is disclosed.

## Supplementary Attachments

**SA1:**
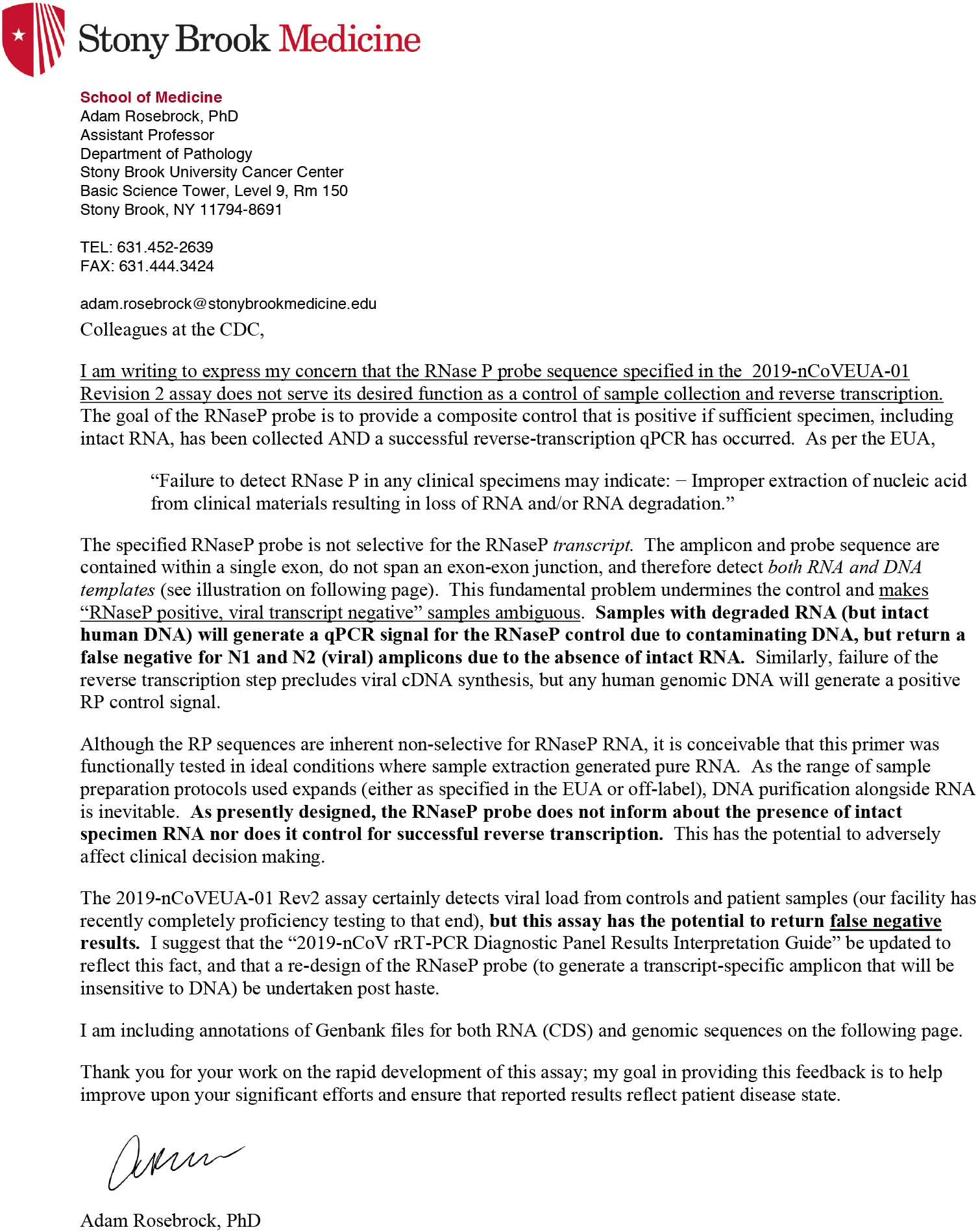
A copy of the author’s letter sent on March 20, 2020 to the CDC feedback line connected with the SARS-nCoV EUA.

**SA2:**
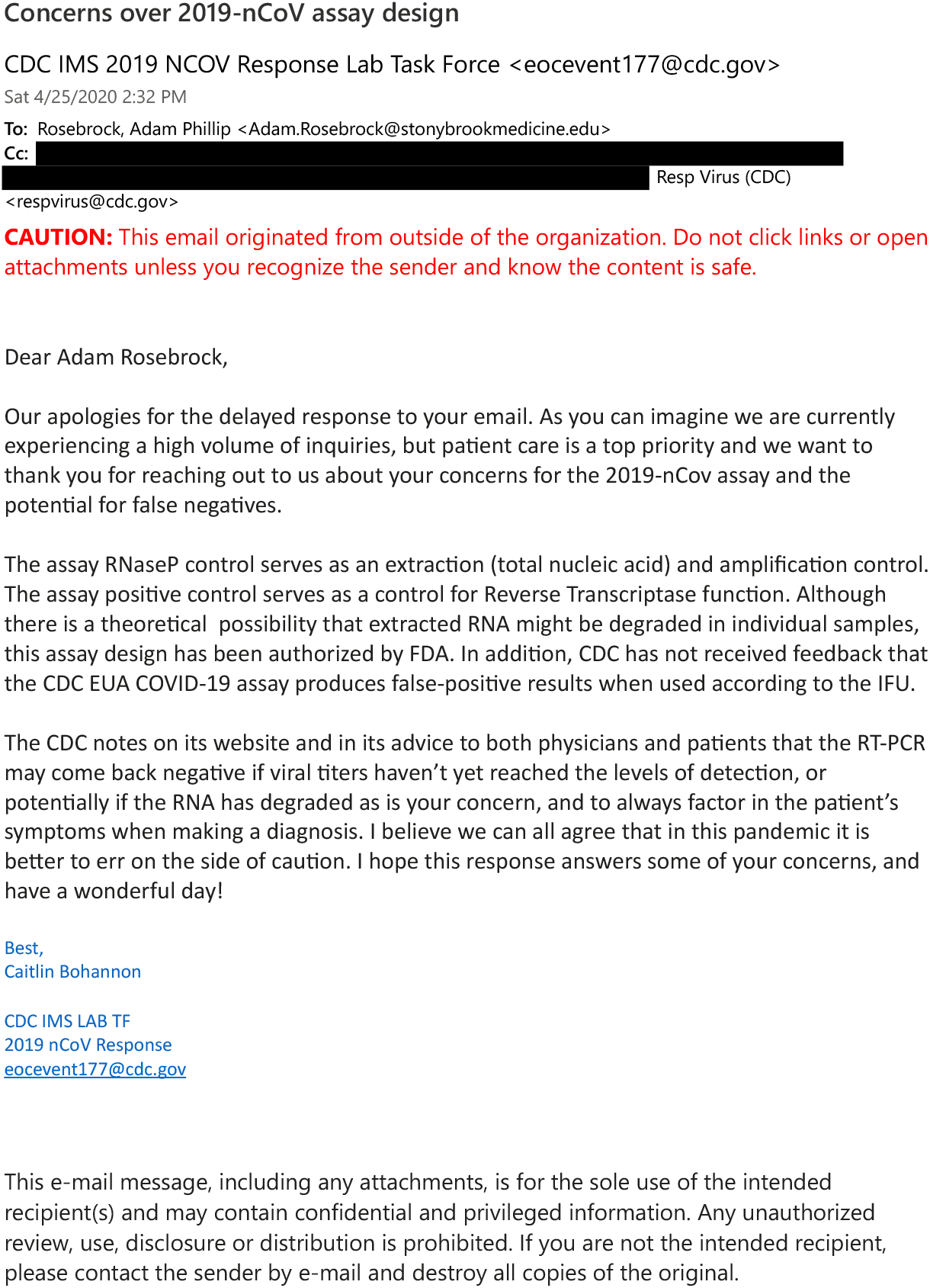
CDC’s response, received April 25, 2020. To protect the email addresses of those carbon copied, CDC-internal email addresses have been redacted. This email is being presented in full and unedited. It is internally ambiguous. The first paragraph correctly notes that my email reflected concerns about false negative potential. This is contradicted in the second paragraph as “false-positive results”, which I hope to be a typo by the CDC. Given the importance of this communication in the context of CDC’s response, I am including a contact-info redacted version of this email in spite of the “do not distribute” message. No patient identifying information is present, and this represents US government work product (from the CDC) and that of a NY state, federally grant funded employee (the recipient).

## Notes

### Competing Interest Statement

The authors have declared no competing interest.

### Summary of Updates

Significant re-formatting of document for submission to peer-review. Supplementary material now contained as a single PDF. Communications with CDC are now properly included.

